# Molecular characterization of SARS-CoV-2 from Bangladesh: Implications in genetic diversity, possible origin of the virus, and functional significance of the mutations

**DOI:** 10.1101/2020.10.12.336099

**Authors:** Md. Marufur Rahman, Shirmin Bintay Kader, S M Shahriar Rizvi

## Abstract

In a try to understand the pathogenesis, evolution and epidemiology of the SARS-CoV-2 virus, scientists from all over the world are tracking its genomic changes in real-time. Genomic studies can be helpful in understanding the disease dynamics. We have downloaded 324 complete and near complete SARS-CoV-2 genomes submitted in GISAID database from Bangladesh which were isolated between 30 March to 7 September, 2020. We then compared these genomes with Wuhan reference sequence and found 4160 mutation events including 2253 missense single nucleotide variations, 38 deletions and 10 insertions. The C>T nucleotide change was most prevalent (41% of all muations) possibly due to selective mutation pressure to reduce CpG sites to evade CpG targeted host immune response. The most frequent mutation that occurred in 98% isolates was 3037C>T which is a synonymous change that almost always accompanied 3 other mutations that include 241C>T, 14408C>T (P323L in RdRp) and 23403A>G (D614G in spike protein). The P323L was reported to increase mutation rate and D614G is associated with increased viral replication and currently most prevalent variant circulating all over the world. We identified multiple missense mutations in B-cell and T-cell predicted epitope regions and/or PCR target regions (including R203K and G204R that occurred in 86% of the isolates) that may impact immunogenicity and/or RT-PCR based diagnosis. Our analysis revealed 5 large deletion events in ORF7a and ORF8 gene products that may be associated with less severity of the disease and increased viral clearance. Our phylogeny analysis identified most of the isolates belonged to the Nextstrain clade 20B (86%) and GISAID clade GR (88%). Most of our isolates shared common ancestors either directly with European countries or jointly with middle eastern countries as well as Australia and India. Interestingly, the 19B clade (GISAID S clade) was unique to Chittagong which was originally prevalent in China. This reveals possible multiple introduction of the virus in Bangladesh via different routes. Hence more genome sequencing and analysis with related clinical data is needed to interpret functional significance and better predict the disease dynamics that may be helpful for policy makers to control the COVID-19 pandemic in Bangladesh.

## Background

The world is suffering from COVID-19, a devastating pandemic caused by a novel coronavirus originating from Wuhan, China(1). Meanwhile, scientists from all over the world are trying to understand the virus better using various processes including genome sequencing. The first reported complete genome sequence was identified in January 3, 2020. (2). More than 141 thousand genomic sequences of SARS-CoV-2 has been submitted in the Global Initiative on Sharing All Influenza Data (GISAID) database (3, 4).

The genomic sequences revealed that the length of the SARS-CoV-2 viral genome is ∼30kb. The longest part of the genome at 5’ end encodes for orf1ab polyprotein whereas the rest of the genome consists of genes for encoding four structural proteins namely surface (S), envelope (E), membrane (M) and nucleocapsid (N), accessory proteins and other non-structural proteins (NSP) encoded by ORF3a, ORF6, ORF7a, ORF7b, ORF8 and ORF10 genes(5). Depending on amino acid changes Forster et. al. reported three central variants (A,B,C) of SARS-CoV-2 where A and C being the most common type in Europe and USA and B being the most common type in East Asia (6). Pachetti el. al. identified multiple mutation hotspot with geographic location specificity. They identified mutations in RNA dependent RNA polymerase (RdRp) gene which is important as some of the proposed drugs are targeting RdRp protein and mutations at in the gene may facilitate the virus to escape from those drugs (7). Yao et. al. identified pathogenic variations depending specific SNVs in the Spike glycoprotein (S) changing viral load and cytopathic effects upto 270 folds (8). Deletions in the viral genomes are also common phenomena and sometimes these are related to severity of the diseases (9–11). Still there is a lack of studies to integrate all the deletions in the whole genome of SARS-CoV-2 globally. This may contribute to understand the pathogenic dynamics of the virus over time. The genetic differences among SARS-CoV-2 strains from different location can be linked with their geographical distributions (12). The recent studies suggested that dry and cold climate can boost the spreading of the infections (13).

The clade and lineage nomenclature is rapidly changing. Specific combinations of 9 genetic markers shows 95% of the hCOV-19 data in GISAID can be further classified in major 6 clades named S, L, V, G, GH, GR (5). Initially the virus was classified into 2, then further into 3 super clades (6, 7). The initial assessment of 3 clades indicates distinct geographic distribution (China, USA and Europe). A team of scientists has developed an open source bioinformatics and visualization toolkit named Nextstrain (www.nextstrain.org) for real-time tracking of pathogen evaluation including SARS-CoV-2 (8). Their clade nomenclature is different but supplementary to Rambaut et al. 2020 (9).

The virus was first reported in Bangladesh on March 8, 2020 as first 3 cases were identified at the Institute of Epidemiology and Disease Research, Dhaka. Currently the country is at the community spread stage and the total number of infected is about 379,738 with 5,555 reported deaths from COVID-19 till 12 October, 2020 (14, 15).

It is important to get more sequences from all over the world which will help us for better understanding of the evolution pattern, disease dynamics, phylogeographic distribution of the clades, design drugs and vaccines etc. In this paper we tried to determine the phylogenetic relationship of Bangladeshi isolates with other isolates from around the world. This can help us to know about the travelling routes of the virus into our country and how it travelled in other parts of the world. We also tried to know the specific mutational differences in our isolates compared to the reference sequence and whether there is any clinico-pathological significance associated with those mutations.

## Methods

### Local sequence retrieval

For retrieval of genome sequences from Bangladesh, we have searched in the GISAID database using search term “Bangladesh” as location. We have downloaded all the relevant sequences from search result in FASTA format and also downloaded patient status metadata, sequencing technology metadata and acknowledgement table separately (accessed on 30 August, 2020).

### Mutation analysis

We have used Genome Detective Coronavirus Typing Tool version 1.13 and CoVsurver enabled by GISAID to analyse our query sequences in FASTA format (16, 17). These tools are free of charge, online based bioinformatic tools that are validated to identify and reassemble novel corona virus isolates. Using these tools, we identified both nucleotide and amino acid mutations and similarities compared to SARS-CoV-2 reference sequence NC_045512 (NCBI) and EPI_ISL_402124 (GISAID).

For functional prediction of mutational changes, we have used two web based tools namely SIFT (Sorting Intolerant From Tolerant) and MutPred2 (18, 19). We also have used USCS SARS-CoV-2 Genome Browser (https://genome.ucsc.edu/cgi-bin/hgGateway?db=wuhCor1) to align mutations along base position and functionally significant areas in the genome (20).(A support vector machine (SVM) based tool namely i-Mutant 3.0 was used for predicting the change of protein stability change and ΔΔ*G* from specific mutations (21).

### Phylogeny analysis

We have used open source bioinformatics visualization platform Nextstrain (nextstrain.org,) for phylogeny analysis of our sequences (22). Pairwise sequence alignment and clade assignment was done using web based Nextstrain tool Nextclade beta version 0.4.9 (23). The GISAID clade identification of the sequences was done using GISAID CoVsurver tool. Further detailed information of different clusters was derived from a preformed interactive web-based tree developed from a subsample of global sequences (∼5000) by neighbour joining method in Nextstrain web interface (https://nextstrain.org/ncov/global, date accessed on 30 August, 2020).

## Results

### Mutation Analysis with functional significance

From GISAID database we have found 329 submissions from Bangladesh (accessed on 30 August, 2020). Out of the 329 submissions, 324 were complete/near genomes. Among the complete genomes, 102 were isolated from female patients, 220 were from male patients and for 2 isolates the gender was not reported. The age range of collected samples were between 8 days to 95 years and the median age was 38 years. The submissions came from a total of 9 laboratories and different sequencing platforms were used by different laboratories. A quality control (QC) analysis was done where 7 sequences were flagged as “bad” (private mutation cut-off was set at 20) and 6 sequences were reported to have more than 5 ambiguous mutations.

Variation analysis from 324 genome isolates revealed a total of 4160 mutation events out of which 4112 are Single Nucleotide Variations (SNVs), 38 deletions (in 30 isolates) and 10 insertions (in 10 isolates). Among the SNVs 2253 were missense mutations in coding regions, 1216 were synonymous mutations and rest were in non-coding part of the genome (Table 1). Among all the SNVs the most common change was C>T (∼41%) and the second most prevalent change was G>A (∼16%). There were 5 large deletions (>50 nucleotide) among which three resulted in deletion of a large portion ORF7a gene and another two deleted the ORF8 gene. The highest number of mutation events (including all SNVs and indels) observed in one isolate (EPI_ISL_445217) was 59 and least was zero as one isolate (EPI_ISL_458133) was identical to the reference genome (NC_045512.3) with 99.4% genome coverage. The average number of mutation events was approximately 13 per isolate.

**Table 1:** Distribution of mutation events along different genomic regions of SARS-CoV-2 Bangladeshi isolates.

| Genome segment* |  | Base Position | Total mutation | Missense Mutation | Synonymous Mutation | Deletion** | Insertion |
| --- | --- | --- | --- | --- | --- | --- | --- |
| Coding region | *ORF1ab* | 266-21555 | 1796 | 1021 | 775 | 2 | 0 |
|  | *S* | 21563-25384 | 528 | 416 | 112 | 3 | 0 |
|  | *ORF3a* | 25393-26220 | 135 | 101 | 34 | 7 | 0 |
|  | *E* | 26245-26472 | 4 | 3 | 1 | 0 | 0 |
|  | *M* | 26523-27191 | 65 | 14 | 51 | 0 | 0 |
|  | *ORF6* | 27202-27387 | 8 | 6 | 2 | 1 | 0 |
|  | *ORF7a* | 27394-27759 | 50 | 9 | 41 | 9 | 0 |
|  | *ORF7b* | 27756-27887 | 8 | 5 | 3 | 0 | 0 |
|  | *ORF8* | 27894-28259 | 70 | 34 | 36 | 2 | 0 |
|  | *N* | 28274-29533 | 949 | 635 | 314 | 0 | 3 |
|  | *ORF10* | 29558-29674 | 13 | 0 | 13 | 0 | 2 |
| Non-coding region | Intergenic | - | 12 | - | - | 2 | 1 |
|  | 5'-UTR | 1-265 | 386 | - | - | 5 | 1 |
|  | 3'-UTR | 29675-29903 | 136 | - | - | 7 | 3 |
|  | **Total** |  | **4160** | **2253** | **1382** | **38\*\*** | **10** |
| *Genes in displayed in italics, **Some deletion events continued over multiple regions |  |  |  |  |  |  |  |

The most common mutation in non-coding region was 241C>T, observed in 96% (312) isolates. The mutation with highest frequency (∼98%) in coding region was 3037C>T (synonymous) and 14408C>T (missense) in ORF1ab gene, and 23403A>G (missense) in S gene. The latter caused D614G amino acid change in the spike protein of the virus. Among the 19 missense SNVs that occurred more than 5 times, 12 were predicted to decrease the stability of their respective protein structure (DDG value less than −0.5kcal/mol) and six of the SNVs were predicted to alter protein function. Among these 19 frequent SNVs, 9 were in T-Cell epitope predicted regions and 5 were in B-Cell epitope predicted regions. The T592I mutation in ORF1ab polyprotein (NSP2) was strongly predicted for CD8+ T-Cell epitope that was also predicted for altered protein function. The P4715L, one of the highest frequent SNVs, occurred in the RNA dependent RNA polymerase (RdRp) region of ORF1ab polyprotein. Beside these, two of the high frequency (86%) SNVs (R203K and G204R) occurred in COVID-19 diagnostic RT-PCR target and B-Cell predicted epitope regions, of which the later was predicted to cause altered function of the nucleocapsid protein (Figure-1). Further analysis revealed two SNVs (L3606F, H15Y) were predicted to cause altered ordered interface and altered transmembrane protein for ORF1ab polyprotein (NSP6) and Membrane (M) protein respectively where L3606F was predicted to cause gain of sulfation at Y360 position of NSP3 protein (Table 2).

**Figure 1:**
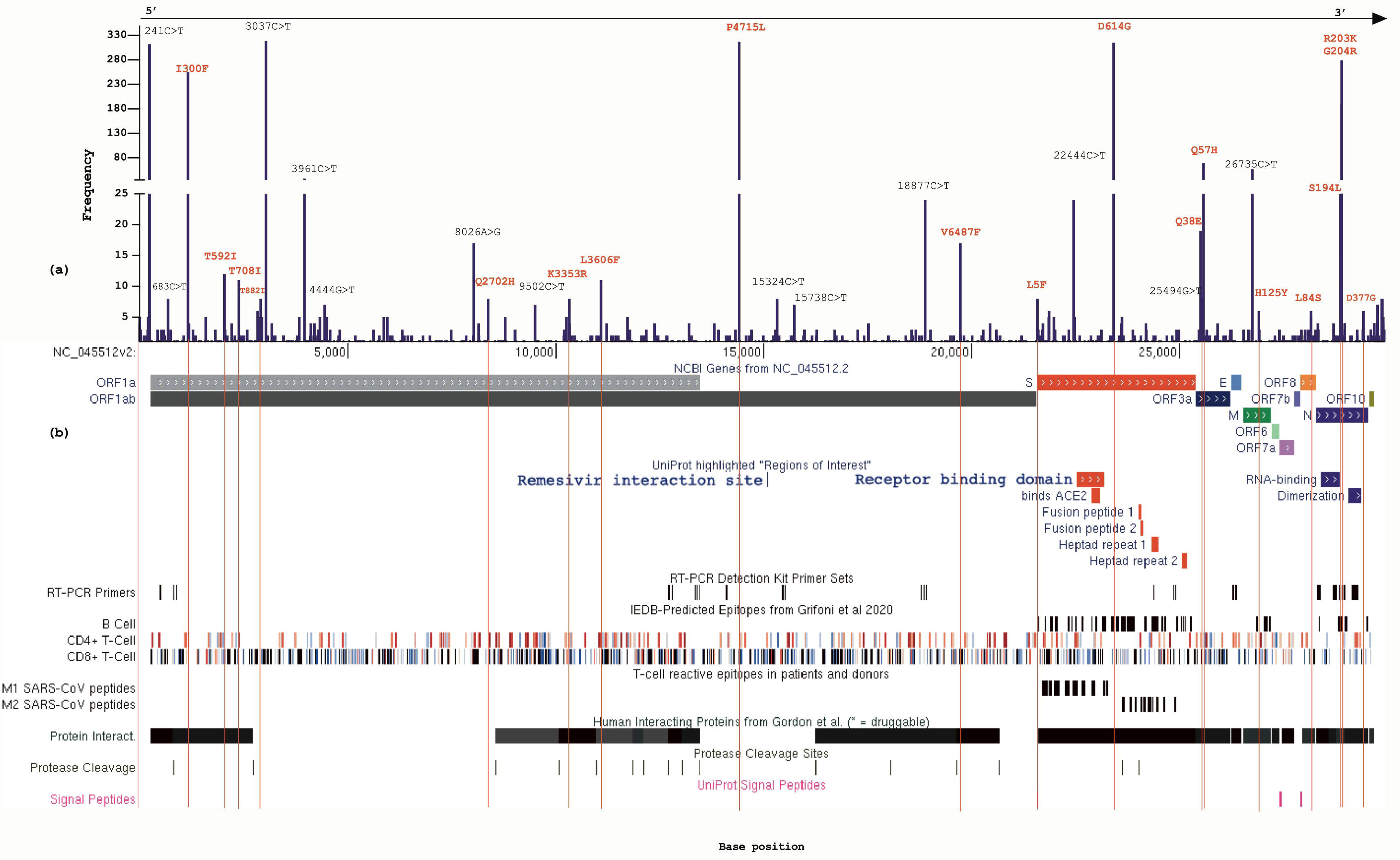
(a) Frequency of SNVs along the base positions of SARS-CoV-2 genome with high frequency (>5) missense SNVs marked red (b) The high frequency missense SNVs aligned (red lines) to different regions of SARS-CoV-2 genome, uniport regions of interest, RT-PCR diagnostic primer set, B-Cell and T-Cell predicted epitope regions, SARS-CoV T Cell epitope regions (M1 and M2 peptides), human protein interaction, protease cleavage and signal peptide regions derived from UCSC genome browser.

**Table 2.** Variants of SARS-CoV-2 genomes observed in more than 5 isolates.

| Genomic change | Type of mutation | Gene | Amino acid change | No. of samples | Structural Prediction Effect (SVM3) | DDG Value (kcal/mol) | Functional prediction Effect | Predicted molecular mechanism change |
| --- | --- | --- | --- | --- | --- | --- | --- | --- |
| 241C > T | Non-coding | 5'-UTR | - | 312 | - | - | - | - |
| 683C>T | Synonymous | ORF1ab | None | 8 | - | - | - | - |
| 1163A > T | Missense | ORF1ab | I300F | 255 | Decrease | -1.79 | Tolerated | CD4+ T Cell epitope |
| 2040C>T | Missense | ORF1ab | T592I | 12 | Neutral | -0.48 | Affect function | CD8+ T Cell epitope (strong prediction) |
| 2388C>T | Missense | ORF1ab | T708I | 11 | Neutral | -0.35 | Tolerated | CD4+ and CD8+ T Cell epitope |
| 2836C>T | Synonymous | ORF1ab | None | 6 | - | - | - | - |
| 2910C>T | Missense | ORF1ab | T882I | 8 | Neutral | -0.10 | Tolerated | - |
| 3037C > T | Synonymous | ORF1ab | None | 318 | - | - | - | - |
| 3961C > T | Synonymous | ORF1ab | None | 38 | - | - | - | - |
| 4444G > T | Synonymous | ORF1ab | None | 7 | - | - | - | - |
| 8026A>G | Synonymous | ORF1ab | None | 17 | - | - | - | - |
| 8371G> T | Missense | ORF1ab | Q2702H | 8 | Decrease | -0.68 | Affect function | - |
| 9502C>T | Synonymous | ORF1ab | None | 7 | - | - | - | - |
| 10323A>G | Missense | ORF1ab | K3353R | 8 | Neutral | -0.13 | Tolerated | CD4+ T Cell epitope |
| 11083G>T | Missense | ORF1ab | L3606F | 11 | Decrease | -1.00 | Affect function | Altered Ordered interface, Altered Transmembrane protein, Gain of Sulfation at Y3607, CD4+ and CD8+ T Cell epitope |
| 14408C> T | Missense | ORF1ab | P4715L | 317 | Decrease | -0.83 | Tolerated | CD8+ T Cell epitope, RdRp zone, |
| 15324C > T | Synonymous | ORF1ab | None | 8 | - | - | - | - |
| 15738C>T | Synonymous | ORF1ab | None | 7 | - | - | - | - |
| 18877C>T | Synonymous | ORF1ab | None | 24 | - | - | - | - |
| 19723G>T | Missense | ORF1ab | V6487F | 17 | Decrease | -1.55 | Tolerated | - |
| 21575C>T | Missense | S | L5F | 8 | Decrease | -0.98 | Tolerated | - |
| 21855C>T | Missense | S | S98F | 6 | Neutral | 0.00 | Tolerated | - |
| 22444C > T | Synonymous | S | None | 24 | - | - | - | - |
| 23403A > G | Missense | S | D614G | 315 | Decrease | -0.93 | Tolerated | B cell epitope |
| 25494G > T | Synonymous | ORF3a | None | 19 | - | - | - | - |
| 25504C>G | Missense | ORF3a | Q38E | 8 | Decrease | -1.02 | Affect function | CD4+ and CD8+ T Cell epitope, transmembrane protein |
| 25563G > T | Missense | ORF3a | Q57H | 30 | Decrease | -2.03 | Affect function | CD4+ T Cell epitope |
| 26735C > T | Synonymous | M | None | 29 | - | - | - | - |
| 26895C>T | Missense | M | H125Y | 6 | Neutral | 0.06 | Tolerated | CD8+ T Cell epitope, Altered Transmembrane protein Altered Ordered interface |
| 28854C > T | Missense | N | S194L | 25 | Neutral | -0.38 | Tolerated | B cell epitope |
| 28881G> A | Missense | N | R203K | 279 | Decrease | -0.93 | Tolerated | B cell epitope, PCR target area, transmembrane protein |
| 28882G>A | Missense | N | R203K | 279 | Decrease | -0.93 | Tolerated | B cell epitope, PCR target area, transmembrane protein |
| 28883G > C | Missense | N | G204R | 279 | Decrease | -0.52 | Affect function | B cell epitope, PCR target area, transmembrane protein |
| 29403A > G | Missense | N | D377G | 6 | Neutral | -0.44 | Tolerated | - |
| 29742G > A | Non-coding | 3'-UTR | - | 7 | - | - | - | - |
| 29848T>A | Non-coding | 3'-UTR | - | 8 | - | - | - | - |
| 29850A>T | Non-coding | 3'-UTR | - | 8 | - | - | - | - |
| 29852A > T | Non-coding | 3'-UTR | - | 7 | - | - | - | - |
| 29853G > A | Non-coding | 3'-UTR | - | 8 | - | - | - | - |

### Mutation analysis of S gene

A separate and detailed analysis of SARS-CoV-2 S gene was done and revealed a total of 530 mutation events among which 414 were missense events and 3 were single amino acid deletions. These mutation events comprised of 56 SNVs and 2 deletions where D614G was the mutation of highest frequency that occurred in 98% (315) isolates. Among the SNVs 35 were predicted to decrease protein stability (DDG less than or around −0.5kcal/mol), 8 were predicted to alter protein function, 18 were in predicted B-Cell epitope region and 26 were in T-Cell predicted epitope region. Individual analysis of these mutations with functional significance is shown in Supplementary Table 1. Three SNVs were found in the receptor binding domain of the spike protein but none of them was in the ACE2 receptor binding part of the protein (Figure 2 and 3).

**Figure 2:**
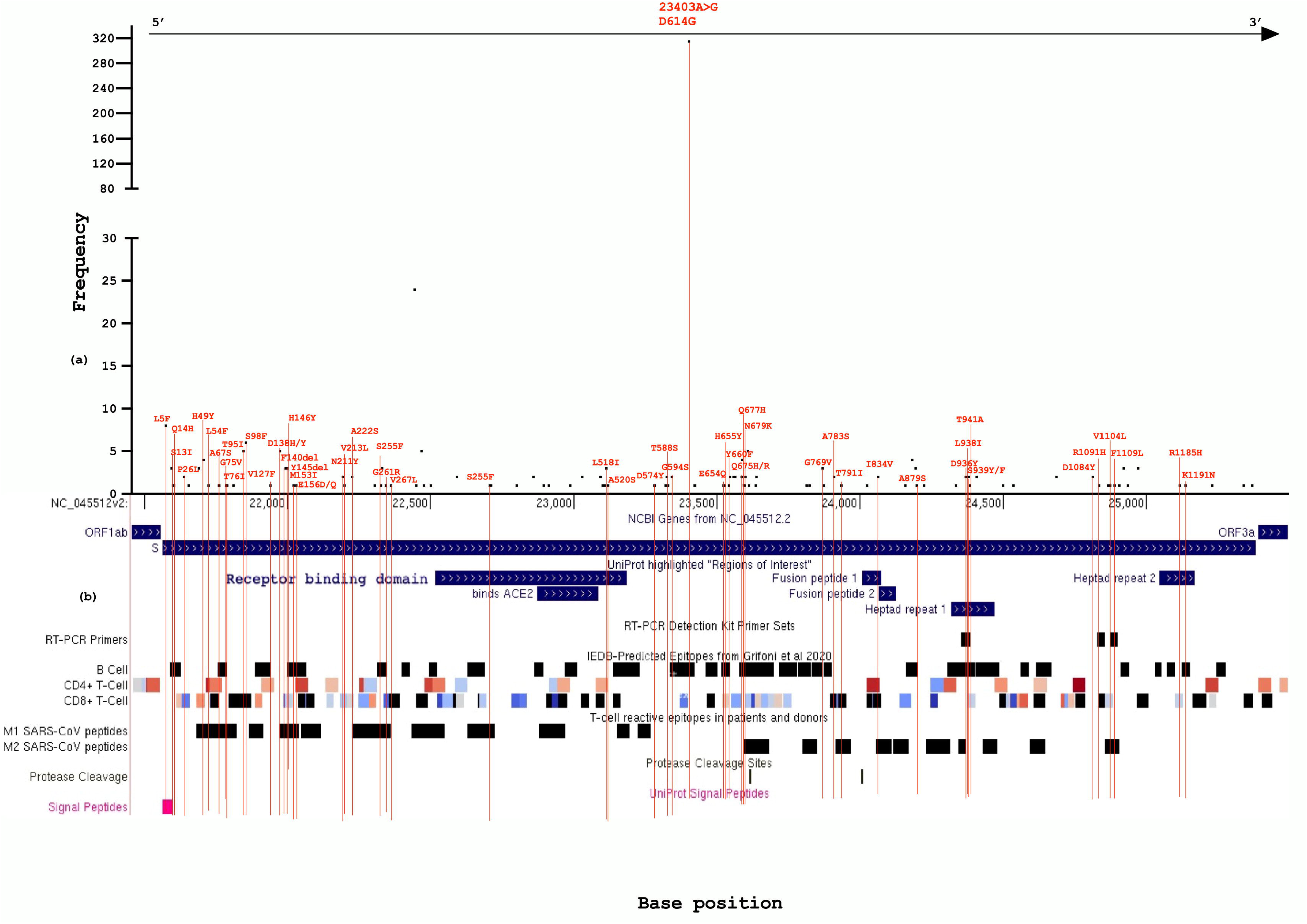
(a) Frequency of SNVs along the base positions of SARS-CoV-2 S gene, missense mutations are annotated in red text (b) The missense SNVs aligned (red lines) to different regions of SARS-CoV-2 S gene, uniport regions of interest, RT-PCR diagnostic primer set, B-Cell and T-Cell predicted epitope regions, SARS-CoV T Cell epitope regions (M1 and M2 peptides), human protein interaction, protease cleavage and signal peptide regions derived from UCSC genome browser.

**Figure 3:**
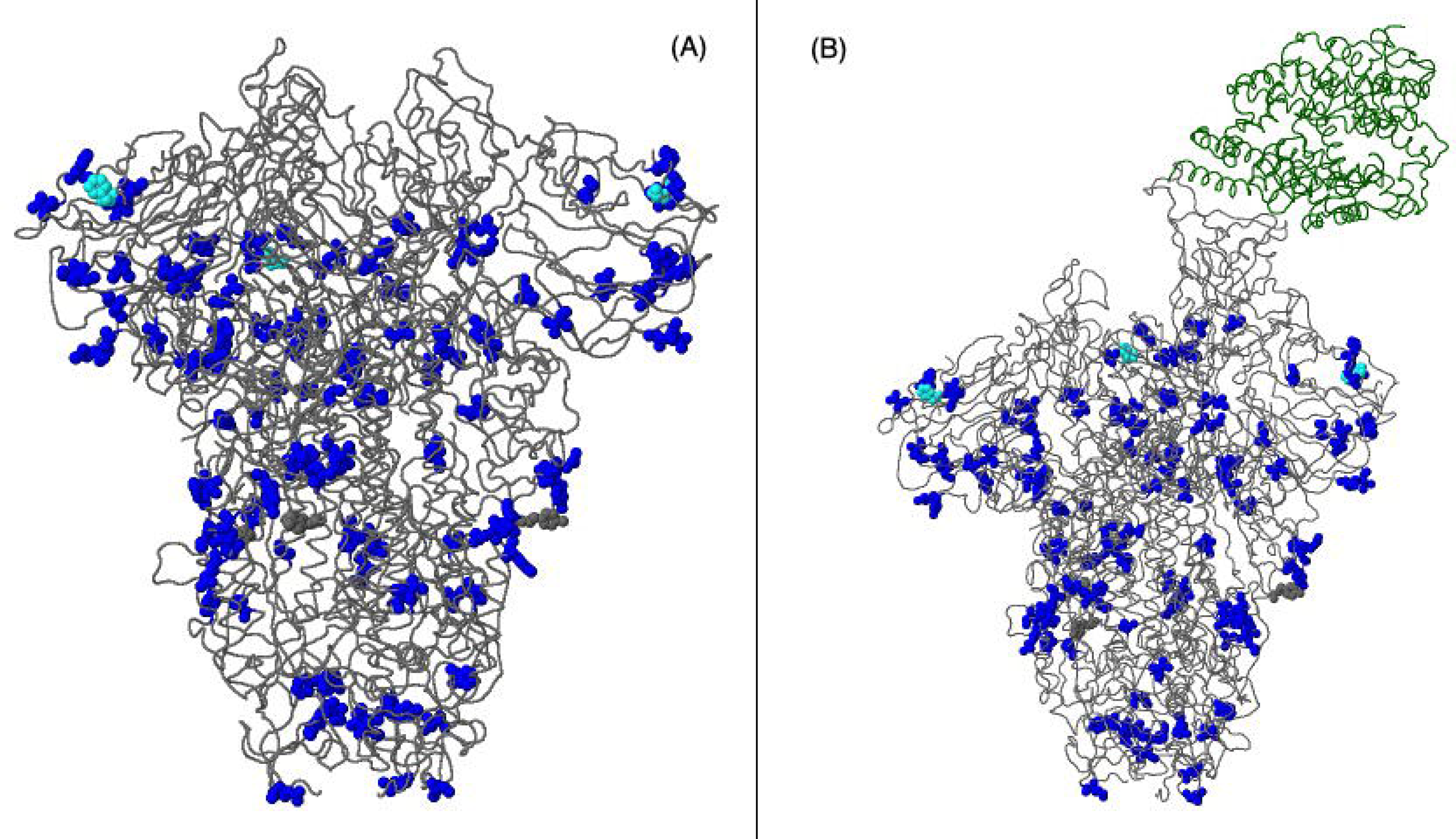
(A) Mutated amino acid positions shown in the unbound structure of SARS-CoV-2 spike protein (except L5F, S13I, Q14H, R1185H and K1195N). (B) Mutated amino acid positions shown in the human ACE2 receptor (marked in green) bound structure of SARS-CoV-2 spike protein (except L5F, S13I, Q14H, R1185H and K1195N).

### Phylogeny analysis

After phylogeny analysis we have found, our 324 isolates were distributed among all the nextstrain clades where 20B clade was the most frequent (86%) (Figure 4). The 19B clade was unique to Chittagong (5 isolates) and one root clade 19A isolate was reported from Dhaka. There were 24 isolates for which the location data was unknown and only one isolate was found in 20C clade. We also have analysed clade distribution of our sequences according to GISAID nomenclature and found more than 96% of the isolates belonged to the G clade and its two major branches GH and GR clade. The common distinctive feature of these three clades is D614G mutation. About 88% of the sequences clustered in GR clade the distinctive feature of which is G204R mutation in the nucleocapsid protein. (Figure 4 and Table 3).

**Figure 4:**
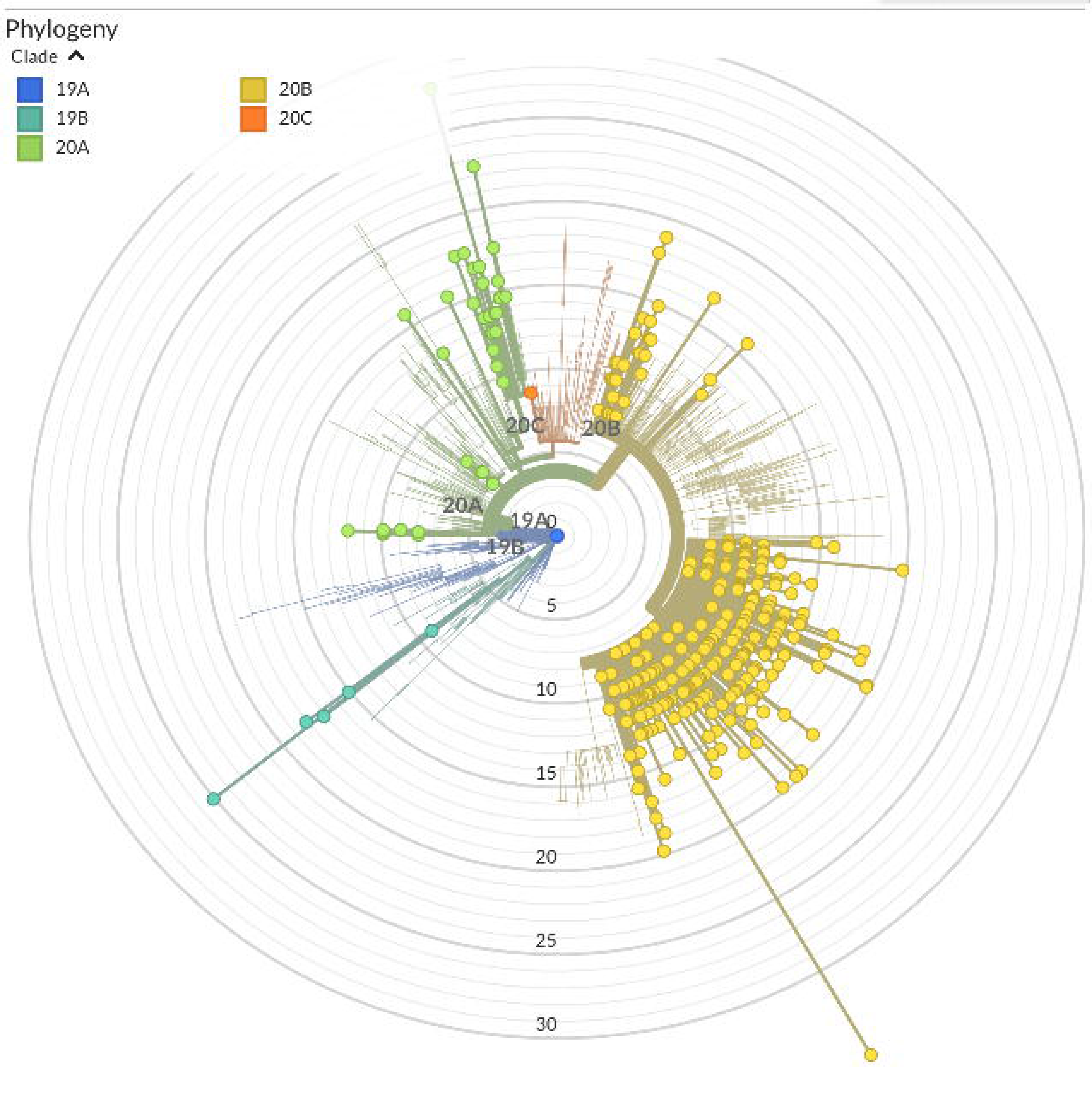
Radial presentation of phylogeny tree formed from 324 SARS-CoV-2 genomes from Bangladesh compared to the reference SARS-CoV-2 sequence shows the isolates belonged to all of the nextstrain clades and the 20B clade is the most prominent.

**Table 3:** Location-wise of distribution of isolates from different clades (24, 25)

| Location (Division) | Nextstrain/GISAID Clade | No. of Isolates | Primary Countries |
| --- | --- | --- | --- |
| Dhaka | 19A/ L | 1/1 | Asia: China/Thailand |
|  | 20A/ G,GH | 6/5,1 | N America/Europe/Asia: USA, Belgium, India |
|  | 20B/ GR,O | 77/75,2 | Europe: UK, Belgium, Sweden |
| Chittagong | 19B/ S | 5/5 | Asia: China |
|  | 20A/ G,GH,O | 14/2,11,1 | N America/Europe/Asia: USA, Belgium, India |
|  | 20B/ GR | 55/55 | Europe: UK, Belgium, Sweden |
| Rajshahi | 20A/ GH | 2/2 | N America/Europe/Asia: USA, Belgium, India |
|  | 20B/ GR | 30/30 | Europe: UK, Belgium, Sweden |
| Rangpur | 20A/ GH | 1/1 | N America/Europe/Asia: USA, Belgium, India |
|  | 20B/ GR,G | 24/23,1 | Europe: UK, Belgium, Sweden |
| Khulna | 20A/ O,GH | 5/3,2 | N America/Europe/Asia: USA, Belgium, India |
|  | 20B/ GR | 22/22 | Europe: UK, Belgium, Sweden |
| Sylhet | 20A/ GH | 2/2 | N America/Europe/Asia: USA, Belgium, India |
|  | 20B/ GR | 19/19 | Europe: UK, Belgium, Sweden |
| Barishal | 20A/ GH | 2/2 | N America/Europe/Asia: USA, Belgium, India |
|  | 20B/ G,GR | 21/1,20 | Europe: UK, Belgium, Sweden |
| Unknown | 20A/G,GH | 6/4,2 | N America/Europe/Asia: USA, Belgium, India |
|  | 20B/GR | 16/16 | Europe: UK, Belgium, Sweden |
|  | 20C/GH | 1/1 | N America: USA |

The overall analysis revealed most of the isolates shared common ancestors with European countries. A subsampled global phylogeny analysis revealed the largest cluster of Bangladeshi isolates shared common ancestor with some Australian isolates that were reported between mid-May to mid-July. Several other clusters were formed sharing common ancestors with countries including Senegal, Morocco, Egypt, Oman, Saudi Arabia, India, Srilanka, Zhenjiang (China), Portugal, Norway, Luxembourg, Bosnia and Herzegovina, England and Italy (Supplementary Figure 1,2,3).

## Discussion

The current SARS-CoV-2 pandemic has changed the world in many ways bringing devastating effects on the society and environment yet we have seen some positive changes and one of which is increased collaboration of scientists and open source projects from all over the world. The collaborative efforts are making huge impact on research and evidence generation. In our study we have tried to gather data on genetic evolution and mutational impacts of SARS-CoV-2 that has been isolated and sequenced in Bangladesh. Our analysis revealed multiple introduction of the virus from different regions in our country as the phylogeny tree shows isolates closely related to different countries and regions of the world. Although most of the isolates were related to isolates from middle eastern and European countries, this can be explained as a lot of Bangladeshi migrant workers live in those countries including Saudi Arabia, Oman, Belgium and Italy. Many of these migrant workers came back to Bangladesh during first and second quarter of this year as number of COVID-19 cases were very high in those regions (26, 27). Interestingly the largest cluster was formed around Australian isolates but going further back on the tree reveals last common ancestor was also related to the European isolates (Sweden and Switzerland). Hence the abundance of 20B clade is observed in Bangladesh unlike neighbouring India and Pakistan where Asian and North American Clades (19A, 19B, 20B) are more dominant (Figure 5).

**Figure 5:**
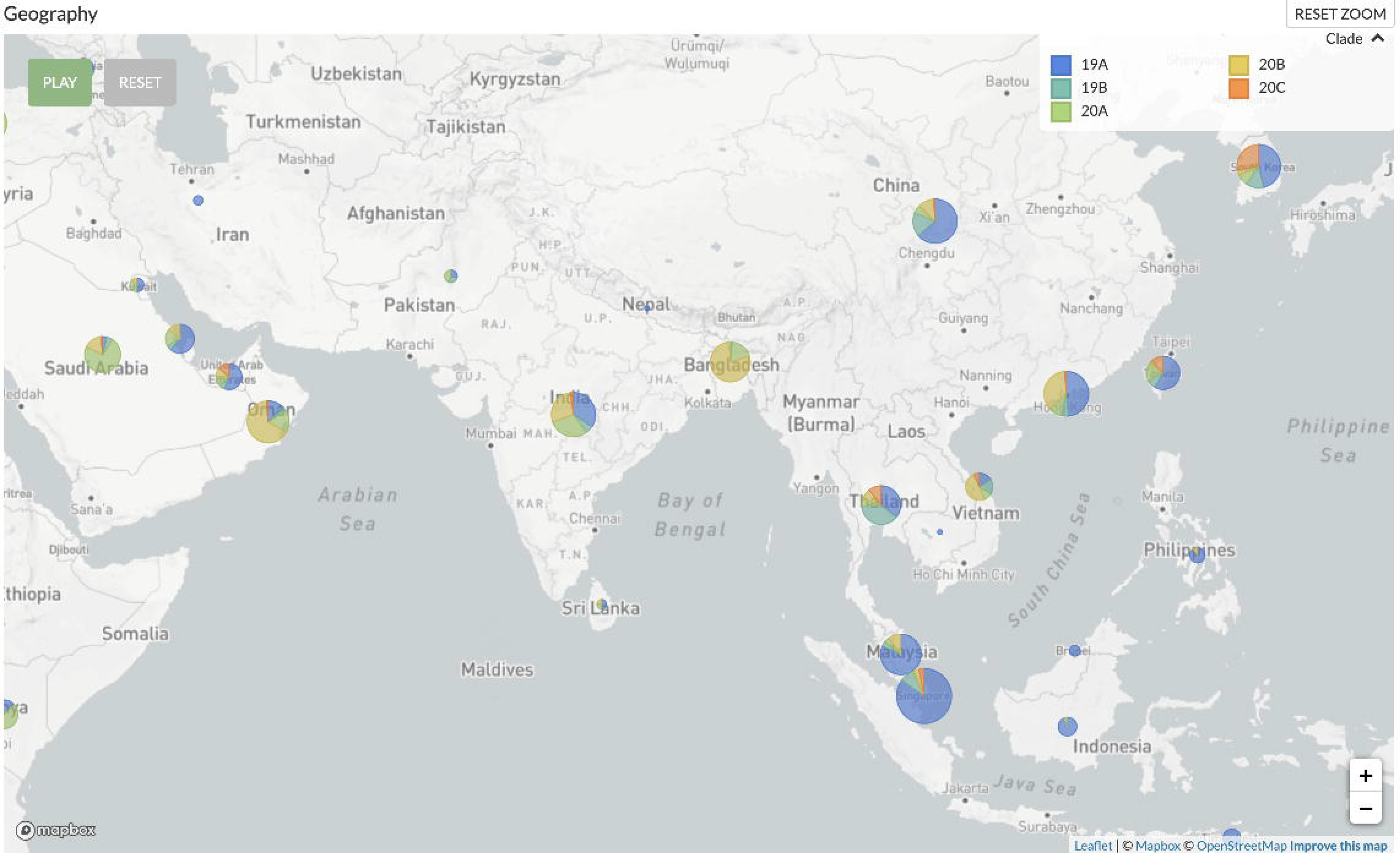
Comparison of clade distribution in different regions of Asia from subsampled global analysis in nextrain.org

In our analysis we have found the most common single nucleotide change was C to T (∼41%). This phenomenon was reported earlier by Biol et al. and can be explained by selective mutation pressure to reduce CpG sites in the presence of abundant human antiviral proteins including APOBEC3 and ZAP. The CpG sites are common targets of viral genome and can be recognized by TLRs that results in release of pro-inflammatory cytokines including type-I interferon, IL-6, IL-12 and TNF-α (28). As pro-inflammatory cytokines play key role in severe COVID-19 including lung tissue damage, the reducing CpG in SARS-CoV-2 may indicate this mutational change facilitates the viral replication but may also reduce disease severity due to less immune reaction through CpG targeted pathways (29). Inverse relation of disease severity with viral load has been reported in a recent study (30). Hence, we think further research is needed focusing the CpG suppression rate in SARS-CoV-2 and its relation with viral load and disease severity which may help designing more potent vaccine and therapeutics. Large deletion events were observed among some of the isolates that resulted in deletion of most of ORF7a or ORF8 gene products. Three such deletions were between base position 27487 to 27552 (65 nucleotide), 27912 to 28256 (344 nucleotide) and 27472 to 27672 (200 nucleotide). Though these proteins are accessory proteins and not necessary for viral replication, ORF7a was found to interact with human ribosomal transport proteins MDN1 and HEATR3 (31). In SARS-CoV the ORF7a was reported to act as cellular translation inhibitor and apoptosis inducer (32). Hence deletion of ORF7a may not change viral replication but can alter disease severity by reducing the chance of ORF7a mediated apoptosis. Similar deletions was reported earlier from USA (33). On the other hand, ORF8 protein is least similar to its SARS-CoV homolog and reported to be associated with MHC-I downregulation that facilitates the virus for immune evasion from Cytotoxic T-Cells. This kind of immune evasion facilitates virus to replicate without getting detected by immune cells hence producing less symptoms. Similar mechanism of immune evasion is observed in some chronic infection causing viruses including HIV-1 and Kaposi Sarcoma associated Herpes Virus (KSHV). This may be one of the reasons for the large number of asymptomatic patients, chronic viral shedding even after clinical cure and redetection of virus long after recovery (34). The ORF8 protein may be responsible for such manifestations and deletion of the protein may cause a disadvantage to the virus increasing more change of getting detected by the immune system and getting eliminated.

The 241C>T mutation was one of the most frequent (96%) one we have observed in our study. This mutation occurred in 5’ UTR region, so it may not have any functional significance except for reducing CpG sites. Interestingly, this mutation always accompanied 3 other mutations in same isolates. Those mutations include 3037C>T (98%), 14408C>T (98%) and 23403A>G (97%) where the latter two are non-synonymous mutations (P4715L in ORF1ab or P323L in RdRp and D614G in Spike protein). This co-occurrence of said mutations are not by chance rather a linkage disequilibrium that has been reported earlier by several studies (35–37). These co-mutations are primary features of GISAID G clade that started rising since February in Europe and now includes more than 70% of all SARS-CoV-2 sequences from all over the world (35). The high prevalence of these 4 mutations in Bangladesh also establishes stronger linkage with European isolates. The P323L mutation in RdRp is reported to be associated with increased mutation rate (38, 39). Among all 4 mutations the D614G is most studied and reported. Though not in the receptor binding part of Spike protein, multiple studies reported that the D614G mutation provides the SARS-CoV-2 an evolutionary advantage for replication. Studies reported D614G mutation increases infectivity of the virus as it was found to be associated with higher viral load and higher infectious titre (40–42).

The D614 amino acid is located between S1 and S2 junction of spike protein. The cleavage of S1-S2 junction by host protease is crucial for entry into the host cell and multiple cleavage site enhances the fusion of SARS-CoV with host cell membrane (43). One study predicted D614G mutation introduces a novel protease (elastase) cleavage site that may enhance the fusion of viral envelop to the host cell membrane hence further facilitate viral RNA entry into the host cell (44). Studies identified the 614G variant of the virus can get more functional advantage in population with delC variant (rs35074065 site) of TMPRSS2 gene (44, 45). This variant is common in Europe, America and South Asia but extremely rare in East Asia according to 1000 Genome project data (46). This may explain the spread of D614G or G clade in mostly Europe, America and recently in South Asia. While some studies suggested that D614G mutation is associated with higher fatality rate (47, 48), several other studies reported no significant association (41, 49). Interestingly one study on US patients reported inverse relation of disease severity and duration with SARS-CoV-2 viral load (30). Considering these analyses it can be said that the current evidence is not clear about the impact of D614G mutation alone on the disease severity and mortality as multiple other stronger factors play role especially age and comorbidity (50).

There has been concerns about the impact of D614G mutation on vaccine development but it is clear that the mutation does not take place in the receptor binding region of the spike protein which is the primary target of the neutralizing antibodies. Also, studies suggested that in natural infection, antibodies generated from D614 variant can cross neutralize G614 variant viruses. Hence this is unlikely that the mutation will have a drastic effect on the immunogenicity of the virus and less like to have any impact on vaccine efficacy (50–52).

The second most frequent mutation in our analysis was a tri-nucleotide change resulting in two amino acid changes which are R203 and G204R in N protein. Our analysis revealed these mutations occurred in a PCR target area and B-cell epitope region. Though functionally tolerated this change was predicted to decrease the protein stability. This finding is similar with other studies that reported R203K and G204K destabilizes N protein structure but may enhance interaction with SARS-CoV-2 Envelop protein that may promote viral release (53, 54).

The I300F mutation (occurred in 78% isolates) was predicted to reduce the stability of NSP2 protein the function of which is not yet confirmed. One study suggested that the amino acid is positioned within the internal groove of the protein and less likely to interact with host factors (55). There two other functionally significant mutations which are Q57H in ORF3a and S194L in N protein. Our analysis suggested the Q57H decreases protein stability with altered protein function may result in loss of a CD4+ T Cell epitope similar to other studies (53, 56). The S194L was predicted to have neutral effect in our study with a reduced DDG value and may attenuate viral assembly as reported in an earlier study (53).

We have observed several mutations in some of the RT-PCR target regions. Though it is yet unknown that if mismatch in primer template changes the accuracy and precision of RT-PCR based COVID-19 diagnosis, we recommend avoidance of using primers containing mutation prone regions for better diagnosis.

As a limitation of our study, we couldn’t derive any clinical information of the patients from whom the samples were collected. The functional significance described in this paper are only computational prediction based and may not always reflect clinical scenario. Also, the genomic sequences were derived using different sequencing platforms (i.e. Illumina, Ion Torrent etc.) and methods (Sanger and Next-generation sequencing) by different laboratories which may have impacted the quality of the sequences hence impacted our analysis. We have found one sequenced that has no mutation compared to the reference sequence which is very unlikely and may possibly be a submission error as the sample was collected long after the original Wuhan outbreak. We hope our findings will create scopes for further research specially including clinical data and also help identifying changes in pathogenicity and infectivity pattern of the virus.

## Supporting information

Supplementary Table 1. Surface glycoprotein variants with their effect and significance.

Supplementary Figure 1: Phylogeny clusters formed by Bangladeshi isolates

Supplementary Figure 2: Phylogeny clusters formed by Bangladeshi isolates

Supplementary Figure 3: Phylogeny clusters formed by Bangladeshi isolates

## Acknowledgements

We acknowledge who were involved in the process of sample collection, genome sequencing and sequence data submission in the GISAID database. We also acknowledge Dr. Senjuti Saha, Scientist, Child Health Research Foundation Bangladesh, who inspired and guided us throughout the writing of this article. This study was funded by the Centre for Medical Biotechnology (CMBT), Management Information System, Directorate General of Health Services, Bangladesh.

## References

1. Zhou P, Yang X Lou, Wang XG, Hu B, Zhang L, Zhang W, et al. A pneumonia outbreak associated with a new coronavirus of probable bat origin. Nature. 2020 Mar 12;579(7798):270–3.

2. Tan W, Zhao X, Ma X, Wang W, Niu P, Xu W, et al. A Novel Coronavirus Genome Identified in a Cluster of Pneumonia Cases — Wuhan, China 2019−2020. China CDC Weekly, 2020, Vol 2, Issue 4, Pages 61–62. 2020 Jan 1;2(4):61–2.

3. GISAID-Initiative. GISAID - Initiative [Internet]. [cited 2020 Sep 19]. Available from: https://www.gisaid.org/

4. Elbe S, Buckland-Merrett G. Data, disease and diplomacy: GISAID’s innovative contribution to global health. Glob Challenges [Internet]. 2017 Jan 1 [cited 2020 May 25];1(1):33–46. Available from: http://doi.wiley.com/10.1002/gch2.1018

5. Khailany RA, Safdar M, Ozaslan M. Genomic characterization of a novel SARS-CoV-2. Gene Reports. 2020 Jun 1;19:100682.

6. Forster P, Forster L, Renfrew C, Forster M. Phylogenetic network analysis of SARS-CoV-2 genomes. Proc Natl Acad Sci. 2020 Apr 8;117(17):202004999.

7. Pachetti M, Marini B, Benedetti F, Giudici F, Mauro E, Storici P, et al. Emerging SARS-CoV-2 mutation hot spots include a novel RNA-dependent-RNA polymerase variant. J Transl Med [Internet]. 2020 Dec 22 [cited 2020 May 9];18(1):179. Available from: https://translational-medicine.biomedcentral.com/articles/10.1186/s12967-020-02344-6

8. Yao H, Lu X, Chen Q, Xu K, Chen Y, Cheng L, et al. Patient-derived mutations impact pathogenicity of SARS-CoV-2. medRxiv. 2020 Apr 23;2020.04.14.20060160.

9. Holmes EC. The Evolution and Emergence of RNA Viruses. [cited 2020 Sep 19]; Available from: www.cdc.gov/eid

10. Armengaud J, Delaunay-Moisan A, Thuret J, Anken E, Acosta-Alvear D, Aragón T, et al. The importance of naturally attenuated SARS-CoV-2 in the fight against COVID-19. Environ Microbiol [Internet]. 2020 Jun 9 [cited 2020 Sep 19];22(6):1997–2000. Available from: https://onlinelibrary.wiley.com/doi/abs/10.1111/1462-2920.15039

11. Pachetti M, Marini B, Benedetti F, Giudici F, Mauro E, Storici P, et al. Emerging SARS-CoV-2 mutation hot spots include a novel RNA-dependent-RNA polymerase variant. J Transl Med [Internet]. 2020 Apr 22 [cited 2020 Aug 8];18(1):179. Available from: https://translational-medicine.biomedcentral.com/articles/10.1186/s12967-020-02344-6

12. Islam MR, Hoque MN, Rahman MS, Alam ASMRU, Akther M, Puspo JA, et al. Genome-wide analysis of SARS-CoV-2 virus strains circulating worldwide implicates heterogeneity. Sci Rep [Internet]. 2020 Dec 1 [cited 2020 Sep 19];10(1):14004. Available from: https://doi.org/10.1038/s41598-020-70812-6

13. Su S, Wong G, Shi W, Liu J, Lai ACK, Zhou J, et al. Epidemiology, Genetic Recombination, and Pathogenesis of Coronaviruses [Internet]. Vol. 24, Trends in Microbiology. Elsevier Ltd; 2016 [cited 2020 Sep 19]. p. 490–502. Available from: https://pubmed.ncbi.nlm.nih.gov/27012512/

14. IEDCR [Internet]. [cited 2020 May 26]. Available from: https://www.iedcr.gov.bd/

15. Reuters. Bangladesh confirms its first three cases of coronavirus - Reuters [Internet]. [cited 2020 May 25]. Available from: https://www.reuters.com/article/us-health-coronavirus-bangladesh-idUSKBN20V0FS

16. Cleemput S, Dumon W, Fonseca V, Abdool Karim W, Giovanetti M, Carlos Alcantara L, et al. Genome Detective Coronavirus Typing Tool for rapid identification and characterization of novel coronavirus genomes.

17. A*STAR-Bioinformatics-Institute. CoVsurver - CoronaVirus Surveillance Server [Internet]. [cited 2020 Sep 19]. Available from: https://corona.bii.a-star.edu.sg/

18. Ng PC, Henikoff S. SIFT: Predicting amino acid changes that affect protein function. Nucleic Acids Res [Internet]. 2003 Jul 1 [cited 2020 Sep 19];31(13):3812–4. Available from: /pmc/articles/PMC168916/?report=abstract

19. Pejaver V, Urresti J, Lugo-Martinez J, Pagel KA, Lin GN, Nam H-J, et al. MutPred2: inferring the molecular and phenotypic impact of amino acid variants. [cited 2020 Sep 19]; Available from: https://doi.org/10.1101/134981

20. Fernandes JD, Hinrichs AS, Clawson H, Gonzalez JN, Lee BT, Nassar LR, et al. The UCSC SARS-CoV-2 Genome Browser [Internet]. Vol. 52, Nature Genetics. Nature Research; 2020 [cited 2020 Oct 11]. p. 991–8. Available from: https://doi.org/10.1038/s41588-020-0697-z.

21. Capriotti E, Fariselli P, Casadio R. I-Mutant2.0: Predicting stability changes upon mutation from the protein sequence or structure. Nucleic Acids Res [Internet]. 2005 Jul [cited 2020 Sep 19];33(SUPPL. 2):W306. Available from: /pmc/articles/PMC1160136/?report=abstract

22. Hadfield J, Megill C, Bell SM, Huddleston J, Potter B, Callender C, et al. Nextstrain: real-time tracking of pathogen evolution. [cited 2020 May 26]; Available from: www.ncbi.nlm.nih.gov

23. Nextclade. Nextclade [Internet]. [cited 2020 Sep 19]. Available from: https://clades.nextstrain.org/

24. GISAID-clade. GISAID - Clade and lineage nomenclature aids in genomic epidemiology of active hCoV-19 viruses [Internet]. [cited 2020 Sep 19]. Available from: https://www.gisaid.org/references/statements-clarifications/clade-and-lineage-nomenclature-aids-in-genomic-epidemiology-of-active-hcov-19-viruses/

25. nextstrain-github.ncov/naming_clades.md at master · nextstrain/ncov · GitHub [Internet]. [cited 2020 Sep 19]. Available from: https://github.com/nextstrain/ncov/blob/master/docs/naming_clades.md

26. Siddiqui T, Sultana M, Sultana R, Akhter S. LABOUR MIGRATION FROM BANGLADESH 2018 ACHIEVEMENTS AND CHALLENGES [Internet]. [cited 2020 Aug 8]. Available from: https://www.forum-asia.org/uploads/wp/2019/05/Migration-Trend-Analysis-2018-RMMRU.pdf

27. IOM. IOM assists vulnerable returning migrants impacted by the COVID-19 pandemic | International Organization for Migration [Internet]. [cited 2020 Aug 8]. Available from: https://bangladesh.iom.int/news/iom-assists-vulnerable-returning-migrants-impacted-covid-19-pandemic

28. Arpaia N, Barton GM. Toll-like receptors: Key players in antiviral immunity [Internet]. Vol. 1, Current Opinion in Virology. Elsevier B.V.; 2011 [cited 2020 Aug 11]. p. 447–54. Available from: /pmc/articles/PMC3311989/?report=abstract

29. Costela-Ruiz VJ, Illescas-Montes R, Puerta-Puerta JM, Ruiz C, Melguizo-Rodríguez L. SARS-CoV-2 infection: The role of cytokines in COVID-19 disease [Internet]. Cytokine and Growth Factor Reviews. Elsevier Ltd; 2020 [cited 2020 Aug 11]. Available from: /pmc/articles/PMC7265853/?report=abstract

30. Argyropoulos K V., Serrano A, Hu J, Black M, Feng X, Shen G, et al. Association of Initial Viral Load in Severe Acute Respiratory Syndrome Coronavirus 2 (SARS-CoV-2) Patients with Outcome and Symptoms. Am J Pathol [Internet]. 2020 Jul [cited 2020 Aug 11];0(0). Available from: https://doi.org/10.1016/j.ajpath.2020.07.001

31. Gordon DE, Jang GM, Bouhaddou M, Xu J, Obernier K, White KM, et al. A SARS-CoV-2 protein interaction map reveals targets for drug repurposing. Nature. 2020 Jul 16;583(7816):459–68.

32. Kopecky-Bromberg SA, Martinez-Sobrido L, Palese P. 7a Protein of Severe Acute Respiratory Syndrome Coronavirus Inhibits Cellular Protein Synthesis and Activates p38 Mitogen-Activated Protein Kinase. J Virol. 2006 Jan 15;80(2):785–93.

33. Addetia A, Xie H, Roychoudhury P, Shrestha L, Loprieno M, Huang ML, et al. Identification of multiple large deletions in ORF7a resulting in in-frame gene fusions in clinical SARS-CoV-2 isolates. Vol. 129, Journal of Clinical Virology. Elsevier B.V.; 2020. p. 104523.

34. Zhang Y, Zhang J, Chen Y, Luo B, Yuan Y, Huang F, et al. The ORF8 Protein of SARS-CoV-2 Mediates Immune Evasion through Potently Downregulating MHC-I. bioRxiv [Internet]. 2020 May 24 [cited 2020 Aug 12];2020.05.24.111823. Available from: https://www.biorxiv.org/content/10.1101/2020.05.24.111823v1

35. Mercatelli D, Giorgi FM. Geographic and Genomic Distribution of SARS-CoV-2 Mutations. Front Microbiol [Internet]. 2020 Jul 22 [cited 2020 Aug 19];11:1800. Available from: https://www.frontiersin.org/article/10.3389/fmicb.2020.01800/full

36. Bai Y, Jiang D, Lon JR, Chen X, Hu M, Lin S, et al. Evolution and molecular characteristics of SARS-CoV-2 genome. bioRxiv [Internet]. 2020 Jun 30 [cited 2020 Aug 19];2020.04.24.058933. Available from: https://doi.org/10.1101/2020.04.24.058933

37. Demir AB, Benvenuto D, Abacioğlu H, Angeletti S, Ciccozzi M. Identification of the nucleotide substitutions in 62 SARS-CoV-2 sequences from Turkey. Turkish J Biol [Internet]. 2020 [cited 2020 Aug 16];44(Special issue 1):178–84. Available from: /pmc/articles/PMC7314507/?report=abstract

38. Begum F, Mukherjee D, Das S, Thagriki D, Tripathi PP, Banerjee AK, et al. Specific mutations in SARS-CoV2 RNA dependent RNA polymerase and helicase alter protein structure, dynamics and thus function: Effect on viral RNA replication. bioRxiv [Internet]. 2020 Apr 27 [cited 2020 Aug 24];2020.04.26.063024. Available from: https://doi.org/10.1101/2020.04.26.063024

39. Eskier D, Karakülah G, Suner A, Oktay Y. RdRp mutations are associated with SARS-CoV-2 genome evolution. PeerJ [Internet]. 2020 Jul 21 [cited 2020 Aug 16];8:e9587. Available from: /pmc/articles/PMC7380272/?report=abstract

40. Korber B, Fischer W, Gnanakaran SG, Yoon H, Theiler J, Abfalterer W, et al. Spike mutation pipeline reveals the emergence of a more transmissible form of SARS-CoV-2. bioRxiv [Internet]. 2020 May 5 [cited 2020 Aug 20];4:2020.04.29.069054. Available from: https://doi.org/10.1101/2020.04.29.069054

41. Korber B, Fischer WM, Gnanakaran S, Yoon H, Theiler J, Abfalterer W, et al. Tracking Changes in SARS-CoV-2 Spike: Evidence that D614G Increases Infectivity of the COVID-19 Virus. Cell [Internet]. 2020 [cited 2020 Aug 20];182:812–827.e19. Available from: https://doi.org/10.1016/j.cell.2020.06.043

42. Zhang L, Jackson CB, Mou H, Ojha A, Rangarajan ES, Izard T, et al. The D614G mutation in the SARS-CoV-2 spike protein reduces S1 shedding and increases infectivity. bioRxiv Prepr Serv Biol [Internet]. 2020 [cited 2020 Aug 22]; Available from: /pmc/articles/PMC7310631/?report=abstract

43. Belouzard S, Chu VC, Whittaker GR. Activation of the SARS coronavirus spike protein via sequential proteolytic cleavage at two distinct sites. Proc Natl Acad Sci U S A [Internet]. 2009 Apr 7 [cited 2020 Aug 20];106(14):5871–6. Available from: www.pnas.org/cgi/content/full/

44. Bhattacharyya C, Das C, Ghosh A, Singh AK, Mukherjee S, Majumder PP, et al. Global Spread of SARS-CoV-2 Subtype with Spike Protein Mutation D614G is Shaped by Human Genomic Variations that Regulate Expression of TMPRSS2 and MX1 Genes. bioRxiv [Internet]. 2020 May 5 [cited 2020 Aug 20];2020.05.04.075911. Available from: https://doi.org/10.1101/2020.05.04.075911

45. Russo R, Andolfo I, Alessandro Lasorsa V, Iolascon A, Capasso M. Genetic analysis of the novel SARS-CoV-2 host receptor TMPRSS2 in different populations. [cited 2020 Aug 20]; Available from: https://doi.org/10.1101/2020.04.23.057190

46. rs35074065 RefSNP Report - dbSNP - NCBI [Internet]. [cited 2020 Aug 20]. Available from: https://www.ncbi.nlm.nih.gov/snp/rs35074065%frequency_tab

47. Toyoshima Y, Nemoto K, Matsumoto S, Nakamura Y, Kiyotani K. SARS-CoV-2 genomic variations associated with mortality rate of COVID-19. J Hum Genet [Internet]. 2020 Jul 22 [cited 2020 Aug 8];1–8. Available from: https://doi.org/10.1038/s10038-020-0808-9

48. Becerra-Flores M, Cardozo T. SARS-CoV-2 viral spike G614 mutation exhibits higher case fatality rate. Int J Clin Pract [Internet]. 2020 Aug 3 [cited 2020 Aug 22];74(8). Available from: https://onlinelibrary.wiley.com/doi/abs/10.1111/ijcp.13525

49. Cassia Wagner, Pavitra Roychoudhury, Chris D. Frazar, Jover Lee, Nicola F. Müller, Louise H. Moncla, et al. Comparing viral load and clinical outcomes in Washington State across D614G substitution in spike protein of SARS-CoV-2 [Internet]. [cited 2020 Aug 22]. Available from: https://github.com/blab/ncov-wa-d614g

50. Grubaugh ND, Hanage WP, Rasmussen AL. Making Sense of Mutation: What D614G Means for the COVID-19 Pandemic Remains Unclear. 2020 [cited 2020 Aug 22]; Available from: https://doi.org/10.1016/j.cell.2020.06.040

51. Ozono S, Zhang Y, Ode H, Seng TT, Imai K, Miyoshi K, et al. Naturally mutated spike proteins of SARS-CoV-2 variants show differential levels of cell entry. bioRxiv [Internet]. 2020 Jun 26 [cited 2020 Aug 22];2020.06.15.151779. Available from: www.gisaid.org

52. Hu J, He CL, Gao Q, Zhang GJ, Cao XX, Long QX, et al. The D614G mutation of SARS-CoV-2 spike protein enhances viral infectivity and decreases neutralization sensitivity to individual convalescent sera. bioRxiv [Internet]. 2020 Jul 6 [cited 2020 Aug 22];2020.06.20.161323. Available from: https://www.biorxiv.org/content/10.1101/2020.06.20.161323v1

53. Wu S, Tian C, Liu P, Guo D, Zheng W, Huang X, et al. Effects of SARS-CoV-2 Mutations on Protein Structures and Intraviral Protein-Protein Interactions. [cited 2020 Aug 24]; Available from: https://doi.org/10.1101/2020.08.15.241349

54. Rahman MS, Rafiul Islam M, Rubayet ASM, Alam U, Islam I, Hoque MN, et al. Evolutionary dynamics of SARS-CoV-2 nucleocapsid protein (N protein) and its consequences [Internet]. bioRxiv. Cold Spring Harbor Laboratory; 2020 Aug [cited 2020 Aug 24]. Available from: https://www.biorxiv.org/content/10.1101/2020.08.05.237339v1

55. Ul Alam ASMR, Rafiul Islam M, Shaminur Rahman M, Islam OK, Anwar Hossain M. Understanding the possible origin and genotyping of first Bangladeshi SARS-CoV-2 strain. J Med Virol [Internet]. 2020 Jun 3 [cited 2020 Aug 24];jmv.26115. Available from: https://onlinelibrary.wiley.com/doi/abs/10.1002/jmv.26115

56. Kim J-S, Jang J-H, Kim J-M, Chung Y-S, Yoo C-K, Han M-G. Genome-Wide Identification and Characterization of Point Mutations in the SARS-CoV-2 Genome. Osong Public Heal Res Perspect [Internet]. 2020 Jun 30 [cited 2020 Aug 24];11(3):101–11. Available from: /pmc/articles/PMC7282418/?report=abstract

