## Supplementary Table 1. Surface glycoprotein variants with their effect and significance. for "Molecular characterization of SARS-CoV-2 from Bangladesh: Implications in genetic diversity, possible origin of the virus, and functional significance of the mutations"

| **Amino acid change** | **No. of isolates** | **Structural Prediction Effect (SVM3)** | **DDG Value (kcal/mol)** | **SIFT prediction** | **Functional Significance** |
| --- | --- | --- | --- | --- | --- |
| L5F | 8 | Decrease | -0.98 | Tolerated | SECReTE motif that facilitates mRNA localization to the ER, present more frequently in (+) ssRNA viruses which utilize ER membranes to create VRCs. Thought to be important for VRC formation by SARS-CoV-2, serves as a possible drug target.  Uniprot Signal peptide region |
| S13I | 1 | Neutral | 0.27 | Tolerated | B-Cell epitope |
| Q14H | 1 | Decrease | -0.77 | Tolerated | B-Cell epitope |
| P26L | 2 | Decrease | -0.69 | Tolerated | CD8+ T-Cell epitope (strong prediction) |
| H49Y | 4 | Neutral | 0.27 | Tolerated | CD8+ T-Cell epitope |
| L54F | 1 | Decrease | -1.14 | Tolerated | CD4+ T-Cell epitope, T cell epitope M1 SARS-CoV peptide mapped to SARS-CoV-2 |
| A67S | 1 | Decrease | -0.60 | Tolerated | CD4+ T-Cell epitope, T cell epitope M1 SARS-CoV peptide mapped to SARS-CoV-2, B-Cell epitope |
| G75V | 1 | Decrease | -0.71 | Tolerated | T cell epitope M1 SARS-CoV peptide mapped to SARS-CoV-2, B-Cell epitope |
| T76I | 1 | Decrease | -0.72 | Tolerated | T cell epitope M1 SARS-CoV peptide mapped to SARS-CoV-2 |
| T95I | 5 | Decrease | -0.78 | Tolerated | CD8+ T-Cell epitope |
| S98F | 6 | Neutral | 0.00 | Tolerated | CD8+ T-Cell epitope |
| V127F | 1 | Decrease | -1.25 | Tolerated | CD4+ T-Cell epitope |
| D138H/Y | 5 | Decrease/Increase | -0.75/-0.07 | Tolerated | (For D138H): Loss of Disulfide linkage at C136, Altered Ordered interface, Loss of Relative solvent accessibility, Altered Transmembrane protein , T cell epitope M1 SARS-CoV peptide mapped to SARS-CoV-2 |
| F140del | 2 | - | - | AFFECT PROTEIN FUNCTION | T cell epitope M1 SARS-CoV peptide mapped to SARS-CoV-2 |
| Y145del | 1 | - | - | Tolerated | CD8+ T-Cell epitope (strong prediction), T cell epitope M1 SARS-CoV peptide mapped to SARS-CoV-2 |
| H146Y | 3 | Increase | -0.02 | Tolerated | CD8+ T-Cell epitope (strong prediction), T cell epitope M1 SARS-CoV peptide mapped to SARS-CoV-2 |
| M153I | 1 | Decrease | -0.98 | Tolerated | B-Cell epitope, T cell epitope M1 SARS-CoV peptide mapped to SARS-CoV-2 |
| E156D/Q | 2 | Decrease | -0.52/-0.98 | Tolerated | - |
| N211Y | 2 | Neutral | 0.13 | AFFECT PROTEIN FUNCTION | Altered Transmembrane protein  CD8+ T-Cell epitope (strong prediction) |
| V213L | 1 | Decrease | -1.23 | Tolerated | CD8+ T-Cell epitope |
| A222S | 2 | Decrease | -0.69 | Tolerated | CD8+ T-Cell epitope, T cell epitope M1 SARS-CoV peptide mapped to SARS-CoV-2 |
| Y248H | 1 | Decrease | -1.20 | Tolerated | CD4+ and CD8+ T-Cell epitope, T cell epitope M1 SARS-CoV peptide mapped to SARS-CoV-2 |
| S255F | 1 | Neutral | -0.03 | Tolerated |  |
| G261R | 1 | Neutral | -0.50 | Tolerated | Altered Ordered interface, Gain of Relative solvent accessibility  Altered Transmembrane protein, Gain of Strand, Gain of ADP-ribosylation at G261, Gain of O-linked glycosylation at S256B-Cell epitope (undetermined prediction score), CD8+ T-Cell epitope (strong prediction), T cell epitope M1 SARS-CoV peptide mapped to SARS-CoV-2 |
| V267L | 1 | Decrease | -0.78 | AFFECT PROTEIN FUNCTION | CD8+ T-Cell epitope |
| V382L | 1 | Decrease | -1.31 | Tolerated | T cell epitope M1 SARS-CoV peptide mapped to SARS-CoV-2 |
| L518I | 3 | Decrease | -1.32 | Tolerated | CD4+ T-Cell epitope |
| A520S | 1 | Decrease | -0.59 | Tolerated | CD4+ T-Cell epitope |
| D574Y | 1 | Neutral | 0.36 | AFFECT PROTEIN FUNCTION | Altered Ordered interface, Loss of Relative solvent accessibility, Gain of Strand, Altered Metal binding, Altered Transmembrane protein, Gain of Sulfation at D574 |
| T588S | 2 | Decrease | -0.44 | Tolerated | - |
| G594S | 2 | Decrease | -1.25 | Tolerated | B-Cell Epitope |
| D614G | 315 | Decrease | -0.93 | Tolerated | Palmitoyltransferase ZDHHC5 and Golgin subfamily A member 7 interaction region, B cell epitope (undetermined prediction score), invasive variant |
| E654Q | 1 | Decrease | -0.67 | Tolerated | CD8+ T-Cell Epitope, B-Cell Epitope |
| H655Y | 1 | Neutral | 0.08 | Tolerated | CD8+ T-Cell Epitope, B-Cell Epitope |
| Y660F | 1 | Neutral | -0.43 | Tolerated | CD8+ T-Cell Epitope, B-Cell Epitope |
| Q675H/R | 6 | Decrease/ Neutral | -0.60/-0.14 | Tolerated | B cell epitope |
| Q677H | 1 | Neutral | -0.67 | Tolerated | B cell epitope |
| N679K | 2 | Decrease | -0.32 | Tolerated | B cell epitope, T-Cell Epitope M2 SARS-CoV peptide mapped to SARS-CoV-2 |
| G769V | 3 | Neutral | -0.49 | Tolerated | Altered Transmembrane protein, B-Cell epitope |
| A783S | 2 | Decrease | -0.75 | Tolerated | B cell epitope |
| T791I | 1 | Neutral | -0.24 | Tolerated | CD8+ T-Cell epitope (strong prediction), T cell epitope M1 SARS-CoV peptide mapped to SARS-CoV-2, fusion peptide in the viral S protein facilitating fusion of viral membrane with host cell membrane, |
| I834V | 2 | Decrease | -0.81 | Tolerated | CD4+ and CD8+ T-Cell epitope, T cell epitope M2 SARS-CoV peptide mapped to SARS-CoV-2, Fusion peptide-1 region |
| A879S | 1 | Decrease | -0.54 | Tolerated | B-Cell Epitope |
| D936Y | 2 | Neutral | -0.35 | AFFECT PROTEIN FUNCTION | B-cell Epitope (undetermined prediction score), Heptad Repeat 1 (HR1), part of virus S protein that combines with HR2 to form a 6-helix bundle that brings viral and cellular membranes into close proximity for fusion and infection, WH1 rt-pcr. Primer region, T cell epitope M2 SARS-CoV peptide mapped to SARS-CoV-2 |
| L938I | 2 | Decrease | -0.87 | Tolerated | B-Cell Epitope, RT-PCR primer region |
| S939Y/F | 2 | Neutral | 0.02 | AFFECT PROTEIN FUNCTION | Heptad Repeat 1 (HR1), part of virus S protein that combines with HR2 to form a 6-helix bundle that brings viral and cellular membranes into close proximity for fusion and infection, WH1 rt-pcr. Primer region, T cell epitope M2 SARS-CoV peptide mapped to SARS-CoV-2, CD8+ T-Cell epitope (strong prediction) |
| T941A | 1 | Decrease | -1.02 | AFFECT PROTEIN FUNCTION | Altered Transmembrane protein, Loss of Relative solvent accessibility, Gain of Helix, Altered Coiled coil  CD8+ T-Cell Epitope, B-Cell Epitope, RT-PCR primer region |
| D1084Y | 2 | Neutral | 0.61 | Tolerated | Altered Metal binding, Loss of Relative solvent accessibility  Gain of Strand, Altered Ordered interface, Gain of Loop, Loss of, Disulfide linkage at C1082, Loss of Catalytic site at C1082 |
| R1091H | 1 | Decrease | -1.36 | Tolerated | WH1 RT-PCR Primer region, T cell epitope M2 SARS-CoV peptide mapped to SARS-CoV-2, CD8+ T-Cell epitope |
| V1104L | 1 | Decrease | -0.70 | Tolerated | CD8+ T-Cell epitope (strong prediction), T cell epitope M2 SARS-CoV peptide mapped to SARS-CoV-2, |
| F1109L | 1 | Decrease | -0.81 | AFFECT PROTEIN FUNCTION | Altered Ordered interface  Altered Disordered interface  Altered DNA binding  Loss of Sulfation at Y1110  T cell epitope M2 SARS-CoV peptide mapped to SARS-CoV-2, NIID rt-pcr Primer region |
| R1185H | 1 | Decrease | -1.57 | Tolerated | CD8+ T-Cell Epitope |
| K1191N | 1 | Decrease | -0.71 | Tolerated | Heptad Repeat 2 (HR2), part of virus S protein that combines with HR1 to form a 6-helix bundle that brings viral and cellular membranes into close proximity for fusion and infection, T cell epitope M2 SARS-CoV peptide mapped to SARS-CoV-2 |
