## Supplementary figures and images for "Molecular characterization of SARS-CoV-2 from Bangladesh: Implications in genetic diversity, possible origin of the virus, and functional significance of the mutations"

### Supplementary Figure 1: Phylogeny clusters formed by Bangladeshi isolates

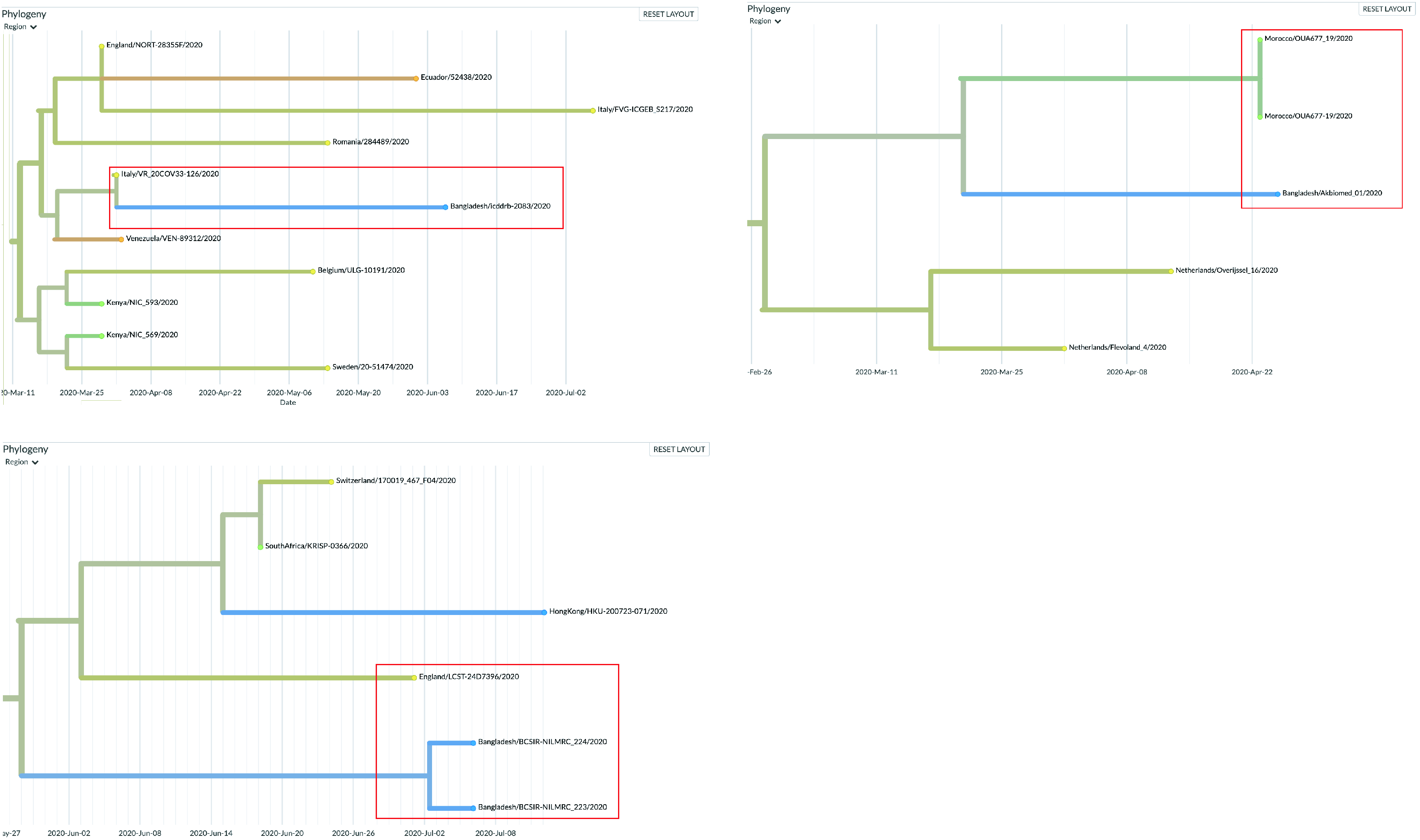

### Supplementary Figure 2: Phylogeny clusters formed by Bangladeshi isolates

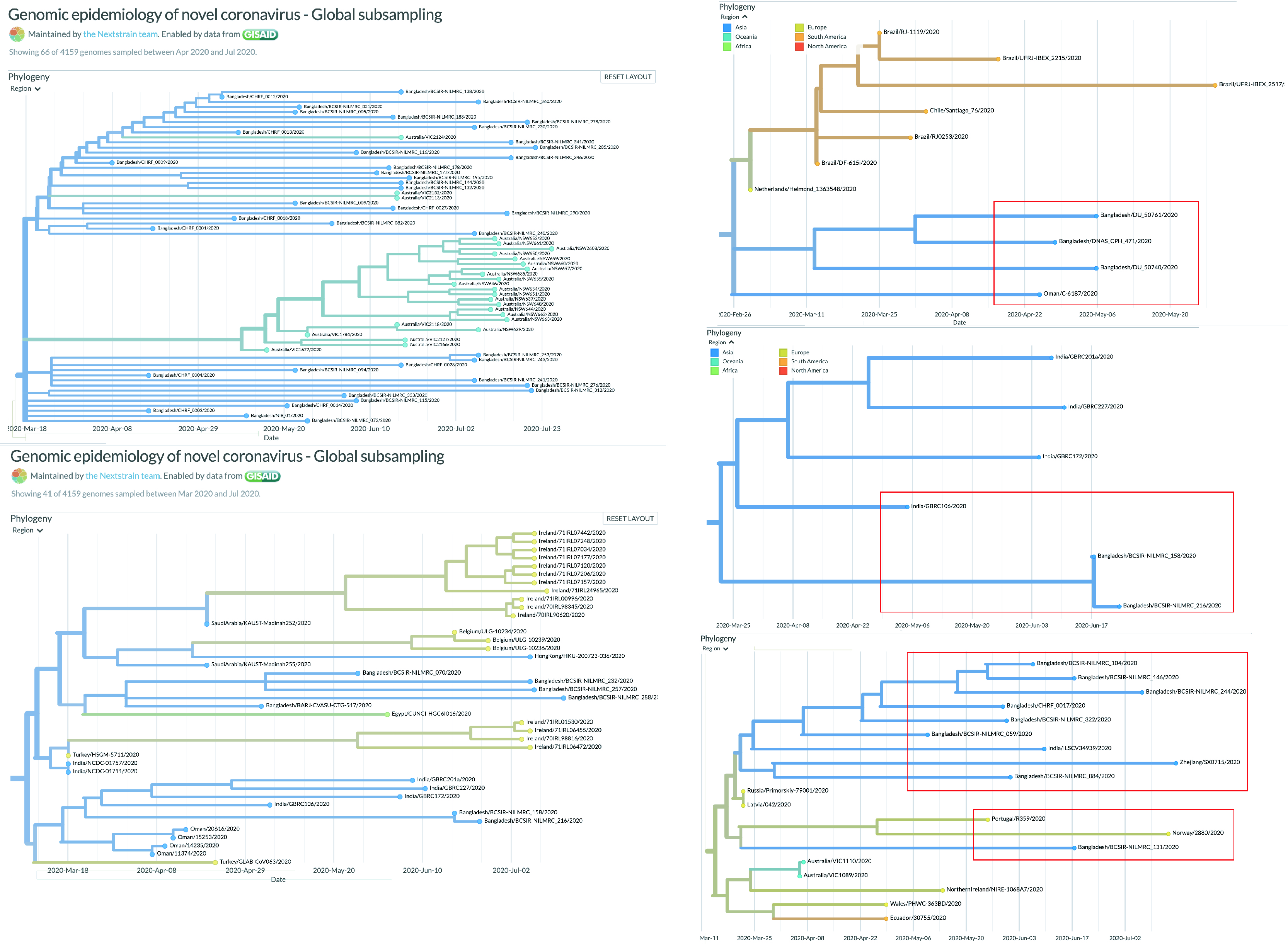

### Supplementary Figure 3: Phylogeny clusters formed by Bangladeshi isolates

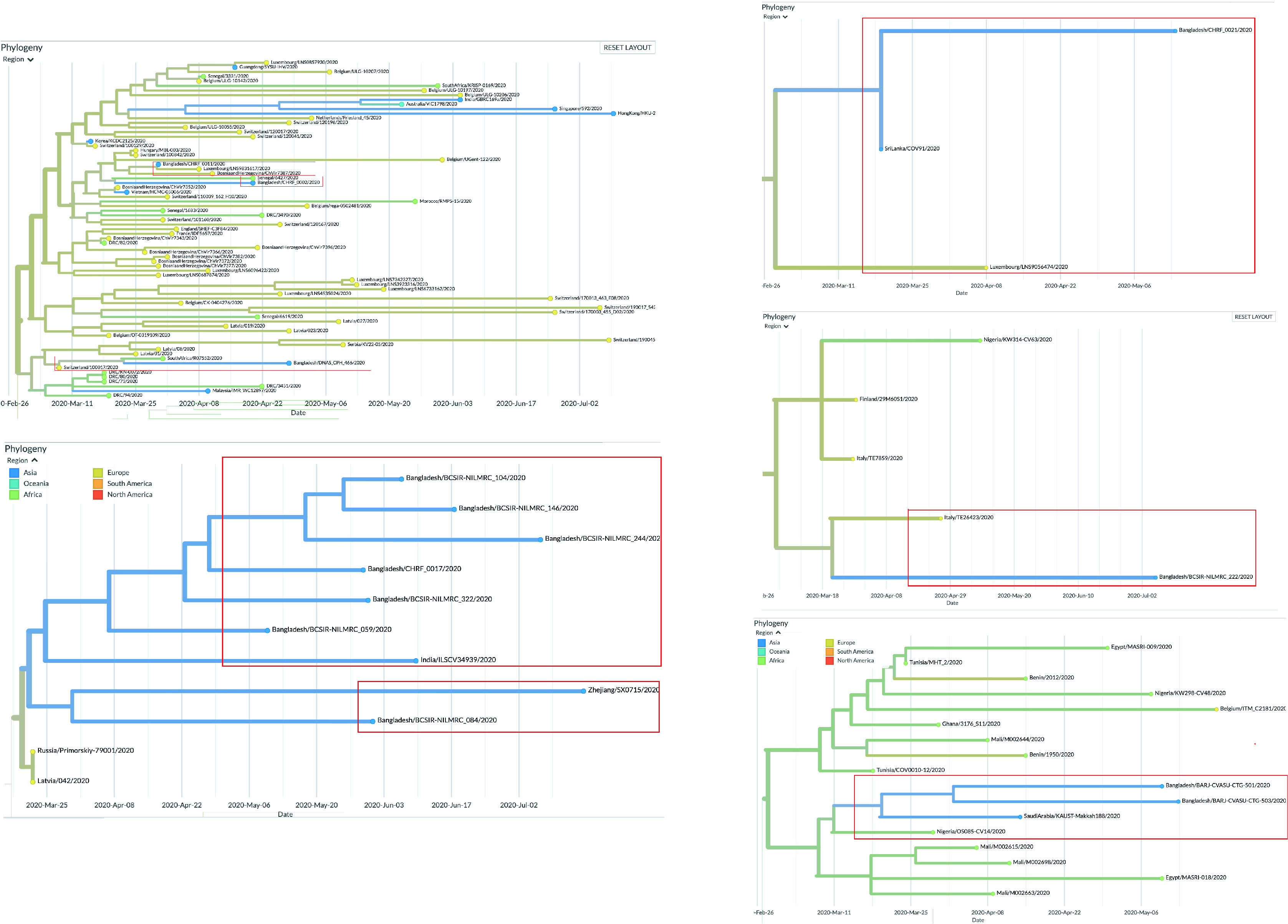
